# Chronic electroconvulsive shock disrupts hippocampal physiology and impairs associative memory in mice

**DOI:** 10.64898/2025.12.24.696412

**Authors:** TR Zhang, NP Vyleta, I Schwein, B Askari, F Vila-Rodriguez, JS Snyder

## Abstract

Electroconvulsive therapy (ECT) is a highly effective treatment for depression but it has undesirable side effects on various aspects of cognition, including memory. Patients might display retrograde amnesia for autobiographical events and anterograde amnesia in tests of recall and recognition memory. These amnestic effects are likely to depend, at least in part, on changes in the hippocampus. ECT-induced volume increases in the hippocampus, and dentate gyrus subregion in particular, correlate with the cognitive deficits. However, the precise cellular and behavioral effects of ECT remain poorly understood. Here, we therefore subjected male and female mice to chronic electroconvulsive shock (ECS) and examined effects on hippocampal physiology and memory. Perforant path inputs onto dentate gyrus neurons were examined using whole cell patch clamp recordings in hippocampal slices. In the intact slice, perforant path stimulation did not result in any synaptic or spiking differences in mice given sham vs ECS treatments. However, GABA_A_ blockade increased depolarization and spiking in sham, but not ECS-treated, mice. Voltage clamp recordings further revealed weaker perforant path excitation of dentate gyrus neurons in ECS-treated mice. Extrapolating these findings to the intact animal, we then used Fos expression as a proxy for neural activity, and found that ECS blunted exploration-induced hippocampal activation, which was most pronounced in the dentate gyrus. Finally, recognition memory tasks revealed that ECS impaired memory for object-context associations but not memory for objects or locations. Collectively, these experiments reveal that a clinically relevant schedule of ECS disrupts synaptic and circuit functions of the hippocampus and causes anterograde amnesia in a test of episodic-like associative memory.

## Introduction

Electroconvulsive therapy (ECT) is the most effective treatment for severe depressive disorder (Group, 2003) but cognitive side effects may deter patients from pursuing it as a treatment option (Koopowitz et al., 2003; Kirov et al., 2016). A major goal is therefore to clarify the neurobiological mechanisms so that procedural refinements can maintain the therapeutic efficacy while minimizing unwanted side effects (Deng et al., 2024).

ECT’s broad effects on cognition are apparent anywhere from hours to months after treatment depending on the task and behavior being tested. Meta analyses indicate cognitive deficits are primarily observed in the days following treatment and that, weeks later, there is recovery and/or improvement (Semkovska and McLoughlin, 2010; Landry et al., 2020). One pattern of initial deficits suggests disruption to the hippocampus and medial temporal lobe systems involved in episodic learning and memory. For example, patients might display retrograde amnesia for autobiographical or public events that occurred prior to treatment (Squire et al., 1975; Lisanby et al., 2000). They also display transient impairments in anterograde recognition memory for discrete items (Squire and Miller, 1974; Steif et al., 1986; Sackeim et al., 2000; Falconer et al., 2010; Fernie et al., 2014). However, recognition memory performance can be ascribed to hippocampal-dependent recollection or neocortex-dependent familiarity judgements (Eichenbaum et al., 2007) though, to date, ECT studies have not differentiated between these 2 processes. Since ECT increases in hippocampal volume are prominent (Oostrom et al., 2018a; Gbyl et al., 2020; Argyelan et al., 2021) and associated with cognition and memory impairments (Oostrom et al., 2018b; Argyelan et al., 2021) one might expect preferential disruption to hippocampal-dependent associative recognition memory over recognition memory for discrete items.

In human studies it is difficult to probe the neurobiological consequences of ECT beyond the level of volumetric changes. Animal models of electroconvulsive shock (ECS) are therefore helpful for identifying how clinically-relevant schedules of ECS impact cellular and circuit functions. Subregional analyses indicate that DG volume increases are particularly robust and are most closely linked to the cognitive deficits (Takamiya et al., 2019; Nuninga et al., 2020; Gbyl et al., 2021). Consistent with volumetric increases in the DG after ECT, ECS in rodents potently upregulates the production of newborn neurons in the adult DG (Madsen et al., 2000; Malberg et al., 2000; Perera et al., 2007; Ueno et al., 2019; Zhang et al., 2021). ECS can reverse long-term potentiation, suggesting it may be capable of disrupting previously-encoded memories (HESSE and TEYLER, 1976). Furthermore, it has been found to increase synaptic strength in the hippocampus and reduce the capacity for subsequent long-term potentiation, aligning with deficits in anterograde learning and memory processes (Stewart et al., 1994; Stewart and Reid, 2000; Chen et al., 2018). Nonetheless, many details remain unknown about how ECS impacts excitatory and inhibitory features of hippocampal circuitry.

To address these gaps, here we subjected mice to chronic ECS and examined DG synaptic transmission and spiking in hippocampal slices, *in vivo* activation (Fos expression) of the hippocampus during learning, and learning and memory in associative and non-associative recognition memory paradigms. These experiments revealed reduced strength of cortical input synapses, blunted expression of the immediate-early gene Fos, and impairments in associative recognition memory. Collectively, these data identify novel, behaviorally-relevant neurobiological consequences of ECS that may be useful for understanding the off-target effects of ECT in humans.

## Methods

### Animals and general experimental design

All procedures were approved by the Animal Care Committee at the University of British Columbia and conducted in accordance with the Canadian Council on Animal Care guidelines. The mice used in this study were Cre-negative animals resulting from a cross between Ascl1^CreERT2^ mice (Kim et al., 2011) and Ai14 mice (Madisen et al., 2010). Mice were weaned at 21 days of age into cages of 2-4 mice where they had *ad libitum* access to food and water and were housed on a 12hr light/dark cycle with lights on at 7am. At 2-3 months old mice were given ECS or sham stimulation. Animals received sham stimulation or ECS every 2 days for a total of 10 sessions over 20 days. Behavior testing began 2 days after the last stimulation session.

### Electroconvulsive shock

Animals were handled daily for one week before ECS to minimize stress associated with manipulation. On each stimulation day, a single animal was removed from its home cage and transferred to an induction chamber containing 5% isoflurane delivered in 20% oxygen. When the animal no longer showed the ability to right itself, anesthesia was reduced and maintained at 2%. The animal was then positioned on a wire platform that supported the body while allowing unrestricted limb movement. Isoflurane (2%) was continuously administered via a nose cone, and a toe-pinch reflex test was performed to confirm adequate anesthesia. After verifying loss of reflexes, both ears were wiped with 70% isopropyl alcohol followed by saline to improve electrical conductivity. Ear clip electrodes were attached, and a conductance test was run to ensure proper contact. ECS was delivered at 30 mA, 100 Hz, with a 0.5-ms square pulse width for a total train duration of 1 s to elicit a tonic-clonic seizure. If an animal showed habituation to stimulation in later sessions, the train duration was increased to 1.2 s and then to 1.5 s as needed; no animal required more than 1.5 s to induce a seizure. After seizure activity had ceased for at least 1 minute, the animal was transferred to a warmed recovery cage and monitored until normal behavior returned, after which it was placed back in its home cage. Sham animals underwent identical procedures except that, following electrode placement, they remained under isoflurane for 2 minutes without receiving any electrical stimulation.

### Behavioral Testing

Mice were habituated to an open field box (40 x 40 x 40cm opaque box) starting 2 days after the last session of ECS. The habituation period consisted of animals being put in the open field arena without any objects for 10 minutes for 2 consecutive days.

On day 3, the animals were trained and tested on a novel object recognition test. In the training phase, animals were put into the familiar open field box with 2 identical objects for 10 minutes. An hour after training, the animals were put back into the box, now with a familiar object and a novel object. The testing phase lasted 5 minutes. Objects were counter-balanced for familiarity and novelty across animals.

On day 4, mice were tested for novel place recognition in the same arena as used to test object recognition. The procedure was the same as novel object recognition except the location, and not identity, of one of the objects changed in the testing phase.

On day 5, mice were tested on novel object-context recognition. Here, mice were placed into two consecutive novel contexts that differed from the one used for object and place recognition in that the walls and floor of the open field box were replaced with materials of different texture and color. New scents were added to the boxes by taping a paper towel with a diluted scent onto the outside walls of the boxes (coffee and vanilla, diluted 1:50 with tap water). In the first context, mice explored a pair of objects (AA) for 10 minutes. After a 3-minute interval in their home cage, mice explored another pair of objects (BB) in the second context. An hour after the second training phase, the animals were put back in the first context and explored 2 objects, one of which was previously seen in that context (A, familiar), and another which had been previously seen but not in that context (B, novel) for 5 minutes.

For behavioral induction of Fos expression, behaviorally naïve mice were allowed to explore the open field box for 10 min, 5 days after chronic ECS treatment. They were then perfused 60 min later and brains were collected for histological processing.

### Tissue Preparation and Immunohistochemistry

Mice were transcardially perfused with 4% paraformaldehyde in phosphate buffered saline and brains were post-fixed for an additional 24 hours. Brains were then transferred to a glycerol-based cryoprotectant solution, and 40 µm coronal sections spanning the hippocampus were cut on an AO860 freezing microtome. For free floating fluorescent immunostaining of c-fos, sections were incubated with 1% H_2_O_2_ for 30min, rabbit anti-Fos antibody (1:1000; Synaptic Systems) for 3 days at 4°C, biotinylated goat anti-rabbit secondary antibody (Jackson Immunoresearch; 1:250) for 1 hour at room temperature, streptavidin-HRP for 1 hour and finally streptavidin-conjugated Alexa 647 (Thermofisher). Slices were then stained with DAPI, mounted onto slides and cover-slipped with PVA-DABCO.

### Imaging and Analyses

All images were acquired with a Leica SP8 confocal microscope. Both dorsal and ventral hippocampal slices were analyzed for all measures. Images of Fos expression were acquired with a 40X oil immersion lens at 1.0 zoom. Images of 1024×1024 pixels (400Hz scan speed) in size at a z-resolution of 1.5µm were merged to encompass the entire hippocampus. Four sections from each animal were chosen (2 dorsal and 2 ventral sections; dorsal Bregma −1.2 to −2.2mm, ventral −2.6 to −3.6mm) and all Fos^+^ cells were counted on ImageJ using the Cell Counter plugin. Cell densities (in mm^2^) were then obtained by dividing cell counts by the analyzed area of the granule cell and pyramidal cell layers.

### Electrophysiology

Within 1 week after the end of a session of chronic ECS, mice were anesthetized with sodium pentobarbital (50 mg/kg, I.P.) and perfused with ice-cold cutting solution containing (in mM): 93 NMDG, 2.5 KCl, 1.2 NaH_2_PO_4_, 30 NaHCO_3_, 20 HEPES, 25 glucose, 5 sodium ascorbate, 3 sodium pyruvate, 10 n-acetyl cysteine, 0.5 CaCl_2_, 10 MgCl_2_ (pH-adjusted to 7.4 with HCl and equilibrated with 95% O_2_ and 5% CO_2_, ∼310 mOsm). Hippocampal slices were cut on a vibratome and transferred to the cutting solution at 35°C for 20 minutes, and then transferred to a storage solution containing (in mM): 87 NaCl, 25 NaHCO_3_, 2.5 KCl, 1.25 NaH_2_PO_4_, 10 glucose, 75 sucrose, 0.5 CaCl_2_, 7 MgCl_2_ (equilibrated with 95% O_2_ and 5% CO_2_, ∼325 mOsm) at 35°C prior to starting experiments.

At ∼32°C, whole-cell patch-clamp recordings were made from unlabelled granule cells that were located in the middle of the granule cell later in the suprapyramidal blade of the dentate gyrus. Slices were superfused with an artificial cerebrospinal fluid (ACSF) containing (in mM): 125 NaCl, 25 NaHCO_3_, 2.5 KCl, 1.25 NaH_2_PO_4_, 25 glucose, 1.2 CaCl_2_, 1 MgCl_2_ (equilibrated with 95% O_2_ and 5% CO_2_, ∼320 mOsm). Recording pipettes were made from 2.0 mm / 1.16 mM (OD/ID) borosilicate glass capillaries and had resistance ∼5 MOhm with an internal solution containing (in mM): 120 K-gluconate, 15 KCl, 2 MgATP, 10 HEPES, 10 EGTA, 0.3 Na_2_GTP, 7 Na_2_-phosphocreatine (pH 7.28 with KOH, ∼300 mOsm). Current-clamp and voltage-clamp recordings were performed at −80 mV. Only recordings with high seal resistance (several gigaohms) and low holding current (less than 50 pA) were included in analyses. For current-clamp recordings, series resistance and pipette capacitance were compensated with the bridge balance and capacitance neutralization circuits of the amplifier.

We first conducted recordings in current clamp to determine intrinsic properties of granule cells (resting membrane potential, input resistance, and rheobase, i.e. minimum current injection needed to elicit spiking). Synaptic and spiking properties were then evaluated in response to medial perforant path stimulation, using trains of stimuli (10Hz, 1sec). Stimuli (0.1 ms) were delivered through a stimulus isolator (A-M Systems analog stimulus isolator model 2200). In ACSF, 10 trains were delivered using stimulus intensities from 100 µA to 1000µA. Synaptic input-output curves were generated by measuring excitatory post-synaptic potential (EPSP) size in response to the first pulse of each train. Spiking was measured over all 10 pulses of each train. The same slices were then superfused with the GABA_A_ blocker, bicuculine methiodide (10 µM), and the same stimulus trains were again delivered to determine the extent to which input-output curves and spiking were constrained by GABAergic inhibition. We then switched to a voltage clamp configuration and measured synaptic currents elicited by medial perforant path stimulation at 100-1000 µA.

### Behavioral analysis

Animal exploration behaviors were manually scored by the experimenter. The frequency and duration of object exploration events were quantified, where an exploration event was defined as direct sniffing or interacting with an object, not including time spent on top of the object looking around the arena.

Recognition memory was assessed by in 2 ways. First, we directly compared the absolute amount of exploration of novel vs familiar stimuli. Second, we calculated novelty-based exploration biases as discrimination indices, defined as (Exploration_NOVEL_ – Exploration_FAMILIAR_) / Exploration_NOVEL+FAMILIAR_. A positive discrimination index indicates that the animal explored the novel object (or object configuration) more often/for a longer period than the familiar object and is interpreted as object memory or preference. A discrimination index of 0 results from equivalent investigation of the 2 objects, indicating a lack of memory or object preference.

### Statistical analyses

All data are provided as supplementary material. Statistical analyses were performed using R and Prism 10 (Graphpad) and graphs were prepared with Prism 10. Sample sizes for each experiment are indicated in the results section. Electrophysiological membrane property data were pooled across sexes since sample sizes were too small to include sex as a factor, and analyzed by t-test. Synaptic transmission and spiking data were fit with linear mixed-effects models with treatment (sham & ECS), sex (male & female), stimulus intensity (100-1000 µA) and drug (ACSF & bicuculline, except the excitatory post-synaptic current (EPSC) experiment, which was conducted entirely in bicuculline) as fixed effects and cell (as appropriate), nested within mouse, as a random effect. The data were then analyzed by ANOVA. Three sham cells and 1 ECS cell were excluded from the EPSP analyses because they spiked in the presence of bicuculline, thereby obscuring the synaptic response. For recognition memory tasks, repeated measures ANOVA was used to analyze effects of ECS on exploration of novel vs familiar stimuli. Discrimination indices were also analyzed by ANOVA (treatment x sex). Two sham-treated mice were excluded from the object recognition analyses because they failed to explore the objects during the tests. In all analyses, significant interactions were followed up with post hoc Holm-Sidak tests. For all analyses, significance was defined as 𝛼 < 0.05.

## Results

### ECS alters perforant path recruitment of DG granule neurons

To test whether ECS caused physiological changes in DG granule cells we performed whole cell patch clamp recordings from neurons located in the middle of the granule cell layer. Intrinsic membrane properties were measured from cells from sham and ECS-treated mice and the sexes were pooled due to limited sample sizes (female sham: 5 cells from 2 mice; male sham: 7 cells from 3 mice; female ECS: 11 cells from 4 mice, male ECS: 11 cells from 4 mice). For all measures, there was no effect of ECS on the membrane properties of DG neurons (Fig. 1A-C): ECS-treated mice had normal resting membrane potential (T_32_=0.3, P=0.8), input resistance (T_32_=0.7, P=0.7) and rheobase, i.e. minimum current injection needed to elicit an action potential (T_32_=1.1, P=0.3).

**Figure 1:**
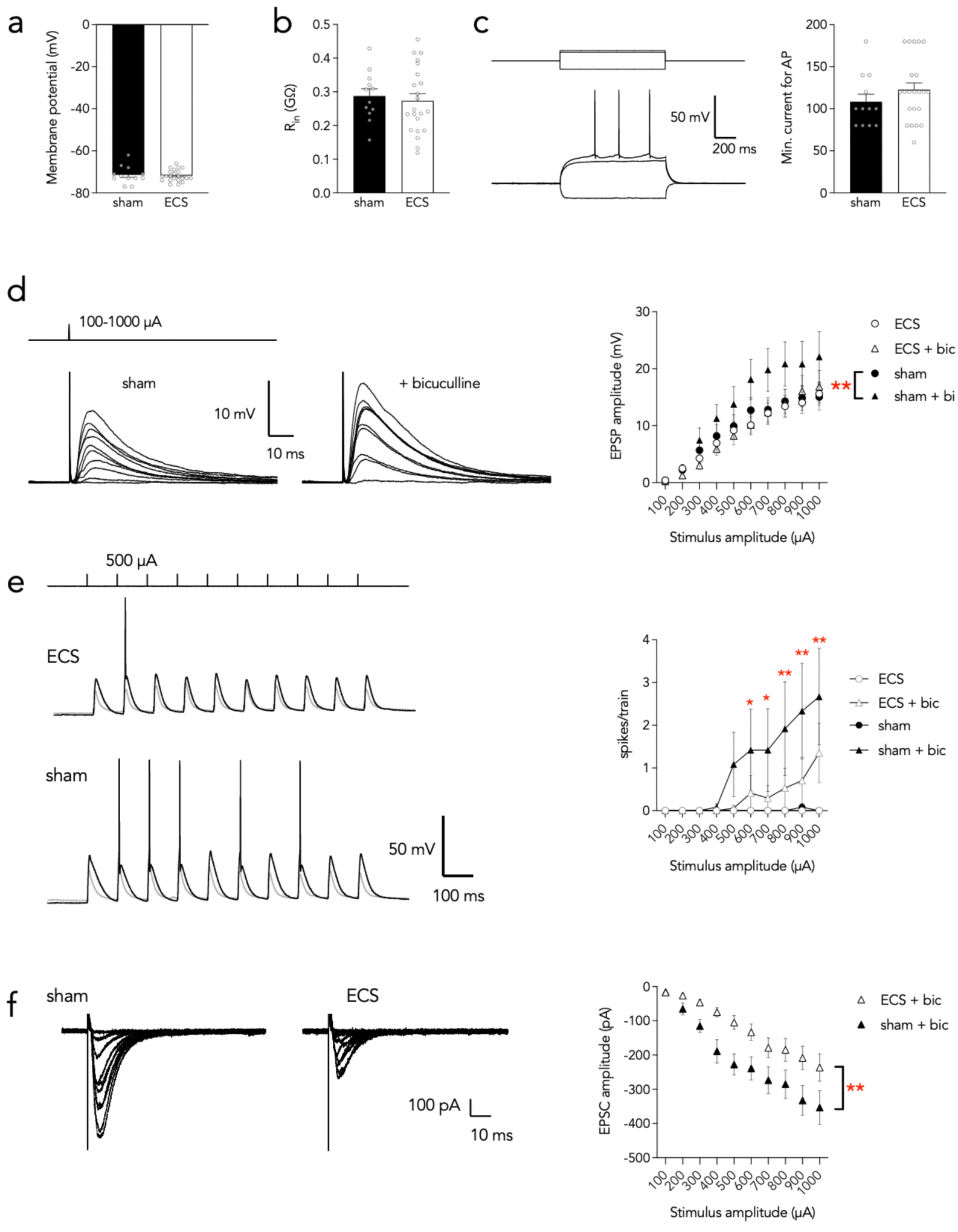
Weaker excitatory synapses and less GABAergic modulation in mice treated with ECS. A) ECS did not alter the resting membrane potential of granule cells. B) ECS did not alter input resistance. C) ECS did not alter the minimum current needed to elicit an action potential. D) GABA_A_ blockade, with bicuculline, increased EPSP magnitude in sham-treated mice but not ECS-treated mice. E) GABA_A_ blockade increased spiking in sham-treated, but not ECS-treated, mice at higher stimulus intensities. Grey traces are voltage recordings (primarily EPSPs) in ACSF and black traces are recordings in bicuculline, which display higher levels of spiking. F) Under GABA_A_ blockade, sham-treated mice had larger EPSCs than ECS-treated mice. Bars and symbols indicate mean ± standard error. *P<0.05, **P<0.01.

We next examined stimulus-response characteristics at medial perforant path inputs in both ACSF and in the presence of bicuculline, a GABA_A_ blocker (cells from sham mice: 5 females, 4 males; ECS mice: 9 females, 7 males). Afferent trains of stimuli (10 pulses at 10Hz, 100-1000 µA) were delivered and EPSPs and postsynaptic spiking were recorded. The magnitude of the first EPSP of the train was used to assess baseline synaptic strength. As expected, synaptic responses increased with greater stimulus amplitude (F_9,29_=4, P=0.001) and in the presence of bicuculline (F_1,7_=10, P=0.02). There was no main effect of ECS treatment (F_1,8_=0.9, P=0.4) or sex (F_1,8_=0.7, P=0.4) but there was an ECS x bicuculline interaction (F_1,7_=11, P=0.01). Post hoc comparisons revealed that EPSPs did not differ between sham- and ECS-treated mice in ACSF but, in the presence of the GABA_A_ blocker bicuculline, EPSP magnitude increased in sham-treated, but not ECS-treated, mice (Fig. 1D; sham mice: ACSF vs bicuculline: P=0.008; ECS mice: ACSF vs bicuculline: P=0.9). All other interactions were not significant (all Ps > 0.06).

Since inhibition profoundly shapes perforant path-DG coupling (Ewell and Jones, 2010; Marín-Burgin et al., 2012), we also measured spiking during the trains of afferent stimulation. As described above, stimulation was performed in ACSF, with GABA intact, and then repeated in the presence of the GABA_A_ blocker, bicuculline (Fig. 1E; cells from sham mice: 5 females, 7 males; ECS mice: 10 females, 7 males). There were no main effects of stimulus intensity (F_9,26_=0.9, P=0.5), bicuculline (F_1,7_=4, P=0.09), ECS treatment (ECS: F_1,13_=4, P=0.08) or sex (F_1,13_=0, P=0.9). Consistent with prior work, mature DG neurons rarely spiked in ACSF but increased spiking during GABA_A_ blockade (averaging ∼1-3 spikes/train; ECS x bicuculline interaction: F_9,425_=10, P<0.0001). Furthermore, a 3-way interaction between stimulus intensity, bicuculline and ECS suggested differences in GABAergic modulation of spiking in sham vs ECS mice (F_9,425_=3, P=0.007; all other interactions P>0.2). We therefore performed pairwise comparisons of spiking in ACSF vs bicuculline, at each of the stimulation intensities, within the sham- and ECS-treated groups. This revealed that GABA_A_ blockade increased spiking in sham-treated, but not ECS-treated, mice at higher stimulus intensities (600-1000 µA; Fig. 1E). Collectively, these results indicate that ECS reduces the GABAergic constraint of spiking in the DG.

To directly probe ECS effects on excitatory transmission, we examined EPSCs in response to medial perforant path stimulation in the presence of bicuculline (Fig. 1F; cells from sham mice: 5 females, 7 males; ECS mice: 10 females, 6 males). As expected, there was a significant effect of stimulation intensity, reflecting the larger EPSCs at higher levels of stimulation (F_9,231_=22, p<0.0001). There were no differences between males and females (F_1,8_=0.3, p=0.6) and no significant interactions involving sex (all Ps > 0.1). However, ECS-treated mice had substantially smaller EPSCs, indicating that ECS directly reduces excitatory synaptic strength in the DG (effect of ECS: F_1,8_=13, p=0.006).

### Chronic ECS reduces exploration-induced Fos expression

We next tested whether chronic ECS has any effects on neuronal recruitment *in vivo*, as measured by expression of the experience-dependent plasticity-related protein Fos (sham mice: 4 males and 4 females; ECS mice: 4 males and 4 females). Following free exploration in an open field, ECS-treated mice had blunted Fos expression in both the dorsal and ventral hippocampus (Fig. 2). While there was no significant interaction between treatment and DG/CA3/CA1 subregions, Fos reduction was most dramatic in the DG (pooled across dorsal and ventral hippocampus, the overall reduction was 59% in DG, 28% in CA3, 6% in CA1). In the dorsal hippocampus, the DG also had the lowest levels of Fos expression (treatment effect: F_1,12_=5.2, P=0.04, subregion effect: F_1.3,15.9_=14, P=0.0009, sex effect: F_1,12_=0, P=0.8; treatment x subregion interaction: F_1.3,15.9_=3, P=0.1; treatment x sex interaction: F_1,12_=0, P=1; treatment x subregion x sex interaction: F_1.3,15.9_=0.4, P=0.6; DG vs CA3, P=0.005; DG vs CA1, P=0.002; CA3 vs CA1, P=0.05). In the ventral hippocampus, in addition to reduced Fos expression in ECS-treated mice, there was also greater Fos in females than in males, and there were differences between all hippocampal subregions (DG < CA3 < CA1; treatment effect: F_1,12_=4.9, P=0.048, subregion effect: F_1.6,18.8_=28, P<0.0001, sex effect: F_1,12_=5.9, P=0.03; treatment x subregion interaction: F_1.6,18.8_=0.1, P=0.9; treatment x sex interaction: F_1,12_=1.6, P=0.2; treatment x subregion x sex interaction: F_1.6,18.8_=0.4, P=0.6; DG vs CA3, P=0.03; DG vs CA1, P<0.0001; CA3 vs CA1, P=0.0004).

**Figure 2:**
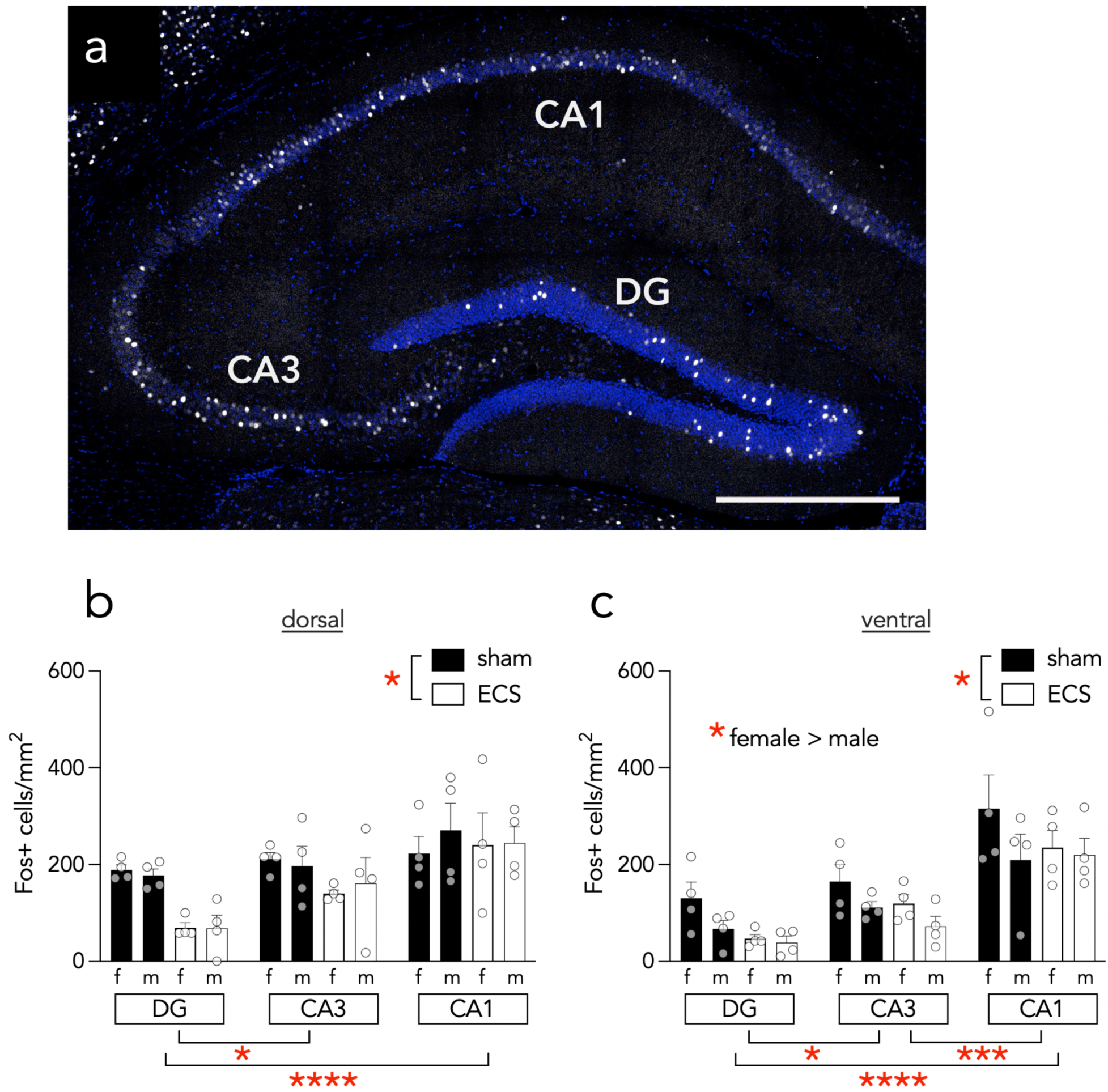
Blunted exploration-induced Fos expression following chronic ECS. A) Representative Fos immunostaining across hippocampal subregions (Fos = white, DAPI = blue, scale bar = 500 µm). B) Five days following chronic ECS, mice showed reduced exploration-induced Fos expression in the dorsal hippocampus compared to sham-treated mice. Fos expression was also lower in the DG. C) ECS-treated mice also showed reduced exploration-induced Fos expression in the ventral hippocampus compared to sham-treated mice. Females has more Fos^+^ cells and all hippocampal subregions differed (DG < CA3 < CA1). Bars indicate mean ± standard error and symbols represent individual mice. *P<0.05, **P<0.01, ***P<0.001, ****P<0.0001.

### ECS impairs associative recognition memory

We finally tested whether ECS-treated mice are impaired on recognition memory tasks that parallel those used to identify anterograde memory deficits in ECT-treated patients (Squire and Miller, 1974; Steif et al., 1986; Falconer et al., 2010; Fernie et al., 2014). Mice were subjected to tests of object, place and object-context recognition memory (sham mice: 10 males, 7 females; ECS mice: 10 males, 5 females. For object recognition 1 sham male was excluded due to a lack of object exploration). Within-subjects analyses of the duration and frequency of exploration events was initially performed by 3-way repeated measures ANOVA (treatment x sex x object novelty). However, since there were no significant main effects of sex or interactions involving sex (all P > 0.05), and to improve the clarity of the figures, we pooled the sexes and performed 2-way (treatment x object novelty) ANOVAs. We have included sex in the discrimination index graphs, however, to show that similar patterns were generally present in males and females. Additionally, the full dataset (i.e. including breakdown by sex) is available as supplementary material.

Overall, mice displayed weak, though statistically significant, object recognition memory (Fig. 3A-E). Mice spent more time exploring the novel object and directed more exploration events towards the novel object (effect of object on duration: F_1,28_=5, P=0.03, effect of object on exploration frequency: F_1,28_=7, P=0.01). ECS-treated mice also displayed generally greater exploration of the objects regardless of whether they were novel or familiar (effect of ECS on duration: F_1,28_=31, P<0.0001, effect of ECS on frequency: F_1,28_=7, P=0.02). However, the treatment x object interactions were not significant, suggesting that ECS does not alter object recognition memory (duration: F_1,28_=3, P=0.09; frequency: F_1,28_=0.7, P=0.4). Analyses of the discrimination indices also revealed that neither ECS nor sex altered object recognition memory (*duration*: effect of sex: F_1,26_=0, P=0.9; effect of ECS: F_1,26_=2, P=0.2; interaction: F_1,26_=3, P=0.1; *frequency*: effect of sex: F_1,26_=0, P=0.8; effect of ECS: F_1,26_=0, P=1; interaction: F_1,26_=0, P=0.9).

**Figure 3:**
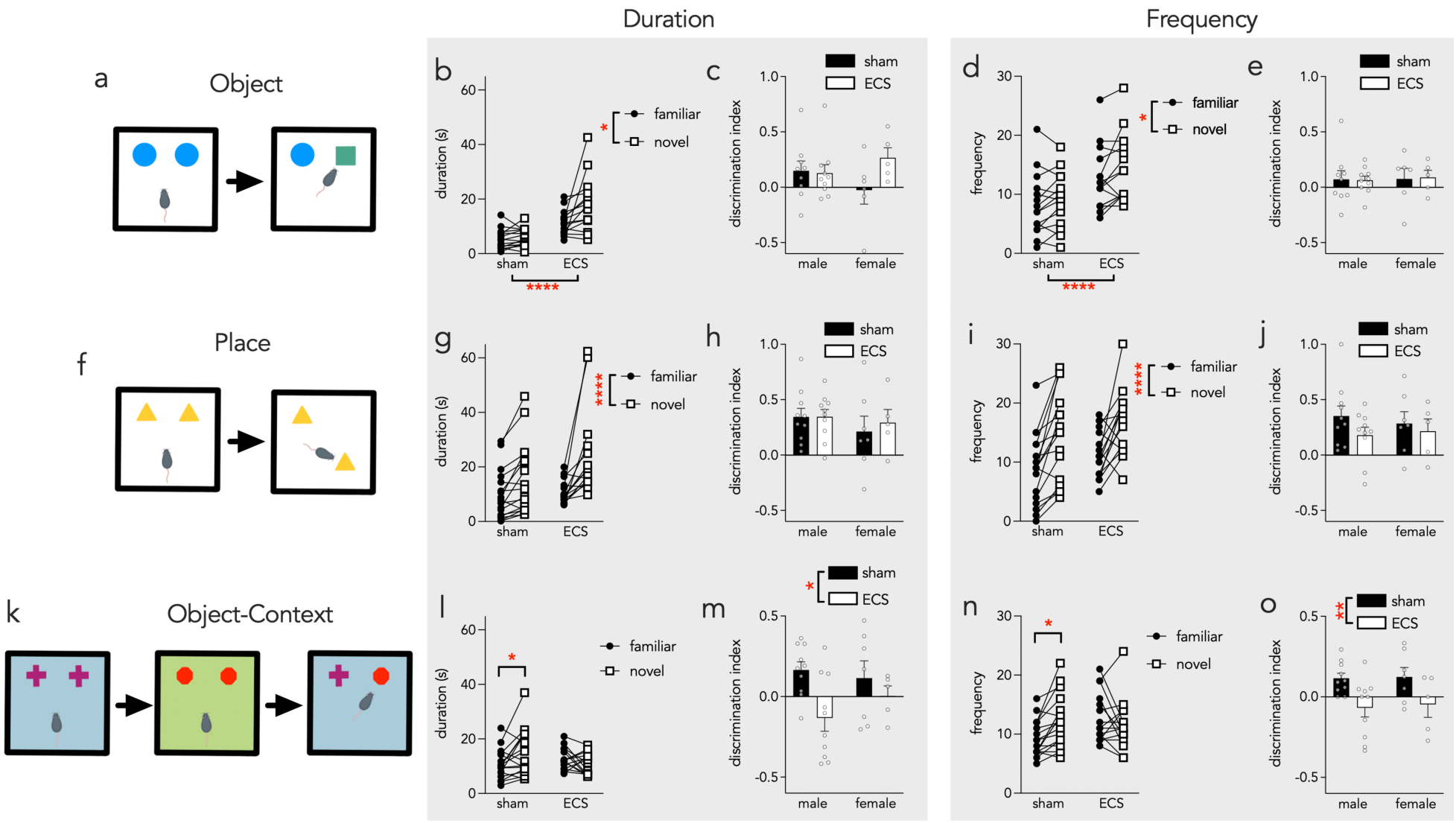
Associative recognition memory is disrupted following ECS. A-E) Object recognition memory. A) Object recognition task schematic. Preferential exploration of a novel object is interpreted as memory. B) At test, ECS-treated mice spent more time exploring objects and, overall, mice spend more time exploring the novel object. C) Duration discrimination indices did not differ between groups. D) Overall, more exploration events were directed towards the novel object and ECS-treated mice displayed more exploration events. E) Sham and ECS-treated mice did not have different discrimination indices. F-J) Place recognition task. Preferential exploration of the novel location is interpreted as place memory. G) Both groups spent more time exploring the novel object location. H) Both groups had comparable, above chance, duration-based discrimination indices. I) Both groups had more exploration events directed towards the novel location. J) Exploration frequency discrimination indices were equivalent between sexes and treatment groups. K-O) Object-context recognition task. Preferential exploration of the novel object-context configuration is interpreted as memory. L) Sham-treated, but not ECS-treated, mice spent more time exploring the novel object-context configuration. M) The sham group’s duration discrimination index was greater than ECS-treated mice. N) Sham-treated mice directed more exploration events towards the novel object-context configuration. O) The sham group’s frequency discrimination index was greater than that of the ECS-treated mice. Bars indicate mean ± standard error. Symbols indicate individual mice. *P<0.05, **P<0.01, ****P<0.0001

In the test of place recognition, sham- and ECS-treated mice showed strong and comparable memory in terms of duration and frequency of exploration events (Fig. 3F-J). Furthermore, sham- and ECS-treated mice spent a comparable time, overall, exploring the objects. (*duration*: effect of object: F_1,30_=24, P<0.0001, effect of ECS: F_1,30_=2, P=0.2, object x ECS interaction: F_1,30_=4, P=0.05; *frequency*: effect of object: F_1,30_=31, P<0.0001, effect of ECS: F_1,30_=1, P=0.3, object x ECS interaction: F_1,30_=0, P=0.8). Discrimination indices revealed a similar pattern, where both groups displayed strong memory and did not differ from one another (*duration*: effect of sex: F_1,28_=0.9, P=0.4; effect of ECS: F_1,28_=0.2, P=0.7; interaction: F_1,28_=0.2, P=0.7; *frequency*: effect of sex: F_1,28_=0, P=0.9; effect of ECS: F_1,28_=2, P=0.2; interaction: F_1,28_=0.3, P=0.6).

Finally, in a test of associative memory, mice were subjected to a test of object-context memory, where they must distinguish a familiar object in a familiar context, but in a novel configuration (i.e. because the object has never been encountered in that context before). Here, sham-treated, but not ECS-treated, mice were able to recognize the novel object-context configuration (Fig. 3K-O). When assessing exploration duration, only sham-treated mice explored the novel configuration more than the familiar one (effect of object: F_1,30_=1, P=0.3, effect of ECS: F_1,30_=0.3, P=0.6, object x ECS interaction: F_1,30_=7, P=0.01; sham familiar vs novel: P=0.02, ECS familiar vs novel: P=0.4). Likewise, analyses of exploration frequency revealed that sham-treated mice selectively recognized the novel object-context configuration (effect of object: F_1,30_=1, P=0.3, effect of ECS: F_1,30_=1.2, P=0.3, object x ECS interaction: F_1,30_=9, P=0.005; sham novel vs familiar: P=0.01, ECS novel vs familiar: P=0.3). Analyses of discrimination indices also revealed selective object-context memory in the sham-treated mice (*duration*: effect of sex: F_1,28_=0.3, P=0.6; effect of ECS: F_1,28_=6, P=0.02; interaction: F_1,28_=1.3, P=0.3; *frequency*: effect of sex: F_1,28_=0, P=0.8; effect of ECS: F_1,28_=10, P=0.004; interaction: F_1,28_=0, P=0.9).

## Discussion

The negative effects of ECT on memory and cognition may depend on changes in hippocampal physiology and plasticity. Here we therefore examined how ECS, in mice, affects neuronal properties that are relevant for understanding its cognitive effects. ECS induced widespread hippocampal hypoactivity, reduced excitatory synaptic strength, and reduced GABAergic control of perforant path-DG coupling. Behaviorally, ECS-treated mice also failed to learn object-context associations, consistent with hippocampal dysfunction (Mumby et al., 2002; Wilson et al., 2013a).

The DG has been implicated in learning since at least as far back as the first report of long-term potentiation in the mammalian brain (Bliss and Lomo, 1973). Since then, evidence has accumulated which links entorhinal-DG/CA3 synaptic plasticity, sparse coding, and excitation-inhibition balance to hippocampal memory processes (Moser et al., 1998; Reagh et al., 2018; Madar et al., 2019; McHugh et al., 2022). Here, we did not observe any changes in intrinsic membrane properties that could influence the excitability of DG neurons, and therefore their recruitment during learning. Instead, ECS reduced the efficacy of perforant path-DG coupling by altering both excitatory and inhibitory aspects of DG circuitry. Using physiologically-relevant trains of stimuli to examine how DG neurons respond to brief episodes of afferent activation, we found that cortical inputs were less effective at inducing spiking. Critically, this effect was dependent on GABAergic systems. With intact inhibition, there were low rates of spiking in both sham and ECS groups, owing to the exceptionally strong GABAergic suppression of DG activity (Coulter and Carlson, 2007; Marín-Burgin et al., 2012). However, only cells from sham-treated mice increased spiking when GABA_A_ receptors were blocked. These results would seem to contrast with a previous report that ECS increases EPSP-spike coupling in the DG (Imoto et al., 2017). And while the direction of the ECS effects may differ, both studies found that ECS-treated mice had reduced GABAergic modulation of spiking, which could critically influence DG pattern separation/completion functions needed for detailed episodic memory formation (Madar et al., 2019; McHugh et al., 2022).

In addition to the effects on spiking, ECS also reduced the strength of perforant path EPSCs, further suggesting that sensory inputs from the cortex may be less efficiently relayed to the hippocampus. This result conflicts with previous findings that perforant path inputs are instead strengthened by ECS, in an LTP-like fashion (Stewart et al., 1994; Stewart and Reid, 2000; Chen et al., 2018). The reason for this discrepancy is unclear but could reflect differences in the species (mouse vs rat) or preparation (*in vivo* vs *in vitro*) used. In any case, the reduced spiking and weaker synaptic transmission is consistent with our *in vivo* finding, as well as a similar report by others (Santiago et al., 2024), that ECS reduces activity in the DG as measured by Fos expression. While sparse codes are thought to be fundamental to DG functions in pattern separation and reducing memory interference, these data suggest that excessive hypoactivity may be detrimental to hippocampal learning and memory.

ECT impairs anterograde learning and memory, as measured by recognition memory tests, where patients study a list of items (e.g. words, images, spatial patterns) and are tested on their ability to recognize novel vs. familiar items at various times after training (Squire and Miller, 1974; Steif et al., 1986; Sackeim et al., 2000; Falconer et al., 2010; Fernie et al., 2014). Generally, recognition memory is most impaired at early timepoints after ECT (minutes to days) and at longer training-testing intervals (hours to days) (Squire and Miller, 1974; Steif et al., 1986). However, some forms of recognition memory, such as spatial recognition, are impaired for months after treatment before recovering (Falconer et al., 2010; Fernie et al., 2014). However, due to the limited number of studies and challenges associated with studying memory in patient populations, it remains unclear exactly which types of memory are impaired and which brain regions are involved. In rodents, recognition memory tests have been adapted to test many aspects of episodic-like memory, including memory for objects, locations, temporal order, and what-when-where-like associations between some or all of these variables (Dere et al., 2007). The hippocampus (and/or afferent entorhinal cortex) is often required for successful performance especially at longer delays (Clark et al., 2000), for associative forms of recognition memory that may be more dependent on hippocampal recollection (Li and Chao, 2008; Sauvage et al., 2008; Langston and Wood, 2010; Wilson et al., 2013a; b; Chao et al., 2015), or when training and test phase stimuli have a high degree of feature similarity (Kirwan et al., 2012; Bekinschtein et al., 2013; Kesner et al., 2015; McHugh et al., 2022). In particular, the DG is important for recognition memory tasks that have a higher degree of discriminative difficulty and are likely to depend on its pattern separation functions (Bakker et al., 2008; Bekinschtein et al., 2013, 2014; Kesner et al., 2015) Our behavioral findings therefore suggest that hippocampal-dependent associative recognition memory may be particularly vulnerable to the anterograde amnestic effects of ECS/ECT since mice were able to remember objects and locations but were unable to associate specific objects with specific contexts.

While this study did not probe causation, our behavioral, electrophysiological and Fos expression data provide evidence that ECS impairs DG circuitry, which could contribute to the anterograde amnesia in associative memory paradigms. Future experiments that prevent ECS/ECT-induced changes in specific brain regions, and further probe the nature of the memory deficits, are needed to identify exactly where and how ECS/ECT alters memory.

